# Isolation and characterization of novel filamentous phages from Swiss-type cheeses infecting the Gram-positive bacterium *Propionibacterium freudenreichii*

**DOI:** 10.64898/2026.07.11.737922

**Authors:** Noël Grosset, Aurélie Nicolas, Julien Jardin, Frank Oechslin, Antoine Culot, Sylvain Moineau, Michel Gautier, Éric Guédon

## Abstract

Filamentous phages infecting Gram-positive bacteria remain largely unexplored. Notably, only two filamentous phages, B5 and Philemon infecting *Propionibacterium freudenreichii*, have been described to date in the phage-rich dairy ecosystem. Although both were genomically characterized, only B5 was confirmed to be an infective filamentous single-stranded DNA phage. The aim of this study was to isolate and characterize new filamentous phages from Swiss-type cheese to investigate their diversity, structural features, host specificity, and potential adaptation to the dairy environment. Thirty raw and pasteurized milk cheeses from France were screened for phages infecting *P. freudenreichii* strains. Eleven phages were isolated, nine of which displayed a filamentous morphology. Named MINOG1 to MINOG9, these filamentous phages exhibited genomic features typical of this morphotype, including small single-stranded DNA genomes with collinear genes organized into functional modules. Comparison with B5 and Philemon revealed sequence divergence ranging from 0.1% to 7%. These phages also exhibited a diverse host range. To further explore phage-*P. freudenreichii* interactions, we screened the genomes of the strains used in this study, as well as additional genomes retrieved from the NCBI database, for CRISPR spacers predicted to target these filamentous phages. Numerous strains contained CRISPR spacers showing 79 to 100% identity to genomic regions of these phages. Two *P. freudenreichii* strains displayed markedly different phage resistance levels despite exact spacer-protospacer matches with phages B5, MINOG1, MINOG2, and MINOG8. Conversely, several strains were resistant to nearly all tested phages despite lacking CRISPR spacers targeting them suggesting the presence of additional defense systems in *P. freudenreichii*.

**IMPORTANCE:** Filamentous phages can play important roles in bacterial ecology by modulating host physiology, population dynamics, and bacterial adaptation to specific environments. However, filamentous phages infecting Gram-positive bacteria remain among the least explored bacterial viruses, and their diversity, ecology, and interactions with their hosts are still poorly understood. This knowledge gap is particularly relevant in dairy ecosystems, where phages are abundant and can influence microbial communities and fermentation processes. In characterizing nine new filamentous phages infecting Propionibacterium freudenreichii from Swiss-type cheeses, this study expands the known diversity of filamentous phages associated with Gram-positive bacteria and provides new insights into phage–host interactions and bacterial defense strategies in dairy-associated bacteria.

## INTRODUCTION

*Propionibacterium freudenreichii* is a Gram-positive non-pathogenic bacterium with high GC content that belongs to the *Propionibacteriaceae* family of the Actinomycetota phylum that also includes the family *Bifidobacteriaceae*, *Brevibacteriaceae,* and *Mycobacteriaceae*. The members of the *Propionibacteriaceae* family were named according to their peculiar metabolic capability, including the production of propionic acid from lactate as a major end product. The bacterial family has recently been divided into three genera: *Cutibacterium*, *Acidipropionibacterium*, and *Propionibacterium* (1). While the *Cutibacterium* genus comprises species found notably on human skin (*e.g*., *Cutibacterium acnes*), the genera *Acidipropionibacterium* and *Propionibacterium* are found in various biotopes (*e.g.*, silage, soil, and human gut), notably in milk and cheese (2, 3). Among dairy-associated *Propionibacterium* species, *P. freudenreichii* is traditionally used as a starter culture in hard and semi-hard Swiss-type cheese, such as Emmental, Comté, Beaufort, and Leerdammer (2, 4). It is mainly responsible for the characteristic round “eyes” and nutty flavor of these cheeses. Eyes formation results from the ability of the bacterium to produce and release CO_2_ during growth, while flavor development arises from the fermentation of lactic acid into propionic and acetic acids, lipolysis of milk fat, and catabolism of amino acids into succinic acid, ammonia, and short chain fatty acids (5–8). Additionally, *P. freudenreichii* can serve as protective cultures due to its production of organic acids, such as propionic acid, which help suppress the growth of spoilage and pathogenic microorganisms. Finaly, beyond its technological role in texture and flavor development as well as antibacterial attributes, some strains have demonstrated probiotic properties both *in vitro* and *in viv*o, including modulation of gut microbiota (9), and anti-inflammatory effects (10–13). *P. freudenreichii* is also capable of producing vitamin B12 (14, 15), a metabolite vital for the formation of healthy red blood cells, nerve function, and DNA synthesis (16).

Bacteriophages (phages) are present in virtually all environments where bacteria are found, and play crucial role in regulating microbiota communities and ecosystems (17, 18). In dairy fermentations, phage infections represent a major technological challenge (19). Virulent phages targeting starter cultures (e.g. *Lactococcus* sp. and *Streptococcus thermophilus*) can interfere with milk fermentation, leading to slow or incomplete acidification, altered texture and flavor, and reduced yield (20). These bacterial viruses persist in the dairy environment and can spread through contaminated milk, equipment, or air (21). To mitigate such issues, strategies include strict hygiene control, rotation or mixing of starter strains, and the use of phage-resistant cultures, particularly those with active defense systems, including CRISPR-Cas systems (21, 22). It should be noted that all dairy lactococcal and streptococcal phages isolated to date have a double-stranded DNA (dsDNA) genome and belong to the *Caudoviricetes* class (tail-containing phages) (23, 24).

Phages infecting *P. freudenreichii* have been mostly isolated from Swiss-type cheese and have a siphophage morphotype with a non-contractile tail and a dsDNA genome of approximately of 50 kb (25–29). However, in 2002, Chopin *et al*. (30) reported the isolation of a filamentous phage, named B5, from a Swiss-type cheese that infects *P. freudenreichii*. This phage was described as the first filamentous phage capable of replicating in a Gram-positive bacterium. Almost two decades later, a second filamentous phage, Philemon, was isolated in Denmark from raw milk Emmental cheese using the dairy-associated strain *P. freudenreichii* PB4 as a host. In contrast to B5, Philemon was characterized exclusively at the genomic level. Earlier, Kim and Blaschek (31) provided the first description of a filamentous phage, CAK1, recovered from the supernatant of the Gram-positive *Clostridium beijerinckii* NCIMB 6444 (formerly *Clostridium acetobutylicum*), but its infectivity and inducibility were not demonstrated. To date, B5, Philemon, and CAK1 remain the only characterized filamentous phages known to be associated with Gram-positive bacteria. However, recent metagenomic analyses of pig fecal samples suggest that filamentous phages targeting Gram-positive hosts may be far more abundant, diverse, and widespread than previously recognized (32).

Filamentous phages belong to the order *Tubulavirales*, which comprises three families: *Inoviridae*, *Paulinoviridae*, and *Plectroviridae* (33). To date, most described filamentous phages, primarily those infecting Gram-negative bacteria, belong to the *Inoviridae* family, whereas filamentous phages infecting Gram-positive bacteria are classified within the *Paulinoviridae* family. Beyond host specificity, filamentous phages share conserved morphological and genetic characteristics (34–36). They produce non-enveloped, flexible filamentous particles ranging from 600 to 2,500 nm in length and 6 to 12 nm in diameter. Their genomes consist of single-stranded DNA (ssDNA) of approximately 5 to 10.5-kb in size, and encoding between 7 to 15 proteins. The archetypal members of the *Inoviridae* family include phages M13, fd, and f1, whereas phages B5 and Philemon represents the prototype of the *Paulinoviridae* family (35).

The aim of this work was to isolate and characterize new filamentous phages that infect the Gram-positive bacterium *P. freudenreichii* in order to explore their diversity, structural features, and evolutionary relationships compared to phages B5 and Philemon. As a starting point, thirty Swiss-type cheese samples made from both pasteurized and raw milk and originating from different manufacturers in France were analyzed. The isolated phages were characterized through host range, transmission electron microscopy, S1 nuclease treatment, next-generation sequencing, bioinformatic analyses, and LC-MS/MS. This comprehensive approach enabled the characterization of nine new filamentous phages infecting a Gram-positive bacterium.

## MATERIALS AND METHODS

### Bacterial strains and culture conditions

Ten *Propionibacterium freudenreichii* strains were kindly provided by either the international microbial resource center CIRM-BIA (INRAE, Rennes, France) (*i.e.*, CIRM-BIA15, CIRM-BIA117, CIRM-BIA129, CIRM-BIA508, CIRM-BIA509, CIRM-BIA1101, CIRM-BIA1102, and CIRM-BIA1402) or the Laboratoires STANDA (Caen, France) (*i.e.*, LSP100 and LSP102). Bacterial strains were grown in yeast extract lactate (YEL) medium (37) and stored at -80°C in the same medium supplemented with 15% glycerol. When plating was required, ten g/L of agar (1%) were added to YEL medium (YELA). Strains were grown in liquid media in microaerophilic conditions in glass tubes with screw cap (two-thirds of volume occupied by liquid medium, one-third by headspace air), at 30°C, without agitation for 3 days. Plates were incubated at 30°C for 4 days under anaerobic conditions in a sealed anaerobic box with Anaerocult A (Merck, Darmstadt, Germany). Growth was monitored spectrophotometrically at 600 nm (OD_600_) and by colony-forming unit (CFU) counting.

### Recovery of *Propionibacterium* phages from cheese samples

Phages infecting *P. freudenreichii* were isolated from 30 cheese samples (**Table 1**). Briefly, 10 g of each cheese were separately mixed with 90 mL of YEL medium and homogenized for 5 min using an ULTRA-TURRAX blender at 20,000 rpm. The mixture was then centrifuged at 3,000 x *g* for 20 min. After filtration through a 0.2 µm Millipore filter, each supernatant was tested for the presence of phages using the ten *P. freudenreichii* strains as potential indicator strains. Two hundreds µL of each exponentially grown bacterial culture (optical density at 600 nm [OD_600_] = 0.4 to 0.7) were incubated with 100 µL filtered cheese supernatant for 15 min at room temperature. Each mixture was then serially diluted, mixed with 3 mL of molten YEL top agar (0.7% agar) and poured onto YELA plates. After 4 days of incubation, the plates were examined, and plaque-forming units (PFU) were enumerated when present (**Table 2**).

**Table 1:** Characteristics of cheeses tested for the presence of *Propionibacterium freudenreichii* phages.

| Cheese sample | Type of cheese | Product | Pressed cooked cheese | Raw milk | Maturing (months) | Origin of manufacture |
| --- | --- | --- | --- | --- | --- | --- |
| A1 | Emmental | Grated | Yes | No | 6 | Finistère* |
| A2 | Emmental | Sliced | Yes | No | 6 | Loire Atlantique* |
| **A3** | **Emmental** | **Grated** | **Yes** | **Yes** | **< 4** | **Jura\*** |
| **A4** | **Emmental** | **Sliced** | **Yes** | **Yes** | **< 3-10** | **Vosges\*** |
| **A5** | **Emmental** | **Grated** | **Yes** | **Yes** | **< 3-10** | **Haute-Marne\*** |
| **A6** | **Emmental** | **Sliced** | **Yes** | **Yes** | **< 4-6** | **Savoie\*** |
| A7 | Emmental | Sliced | Yes | Yes | 3 | Ain* |
| A8 | Emmental | Sliced | Yes | No | < 4 | Mayenne* |
| A9 | Emmental | Sliced | Yes | Yes | < 4 | Loire Atlantique* |
| **A10** | **Emmental** | **sliced** | **Yes** | **Yes** | **3-4** | **Haute Saône\*** |
| A11 | Emmental | Sliced | Yes | Yes | 3-4 | Haute Marne* |
| A12 | Emmental | Sliced | Yes | No | < 4-6 | Pas de calais* |
| A13 | Emmental | Sliced | Yes | No | < 4 | Vosges* |
| A14 | Emmental | Sliced | Yes | Yes | < 4 | Savoie* |
| A15 | Comté | Sliced | Yes | Yes | < 4-6 | Jura* |
| **A16** | **Comté** | **Sliced** | **Yes** | **Yes** | **6** | **Doubs\*** |
| A17 | Comté | Sliced | Yes | Yes | 12 | Jura* |
| A18 | Comté | Sliced | Yes | Yes | < 6 | Doubs* |
| **A19** | **Comté** | **Sliced** | **Yes** | **Yes** | **< 6** | **Doubs\*** |
| A20 | Comté | Sliced | Yes | Yes | < 6 | Jura* |
| A21 | Comté | Sliced | Yes | Yes | < 12 | Jura* |
| **A22** | **Comté** | **Sliced** | **Yes** | **Yes** | **< 5** | **Doubs\*** |
| **A23** | **Comté** | **Sliced** | **Yes** | **Yes** | **12** | **Jura\*** |
| A24 | Comté | Sliced | Yes | Yes | 18 | Jura* |
| A25 | Comté | Sliced | Yes | Yes | 24 | Jura* |
| A26 | Leerdammer | Sliced | Yes | No | 6 | Netherlands |
| A27 | Saint-Nectaire | Sliced | No | Yes | 8 | Puy de Dôme* |
| A28 | Beaufort | Sliced | Yes | Yes | 12 | Savoie* |
| A29 | L'Etivaz | Sliced | Yes | Yes | 5 | Switzerland |
| A30 | Ossau | Sliced | No | Yes | 6 | Pyrénées atlantiques* |

\* Department of France. Bold, indicates phage-positive sample (see Table 2).

**Table 2.** ***Propionibacterium freudenreichii* phages titers in various cheese samples.**

| Cheese sample* | Titer of phages (PFU per gram of cheese) on indicator strains |  |  |  |  |  |
| --- | --- | --- | --- | --- | --- | --- |
|  | CIRM-BIA15 | CIRM-BIA129 | CIRM-BIA1101 | CIRM-BIA509 | LSP100 | LSP102 |
| A3 | $1.2 \times 10^3$ | 0 | 0 | $1.2 \times 10^3$ | 0 | 0 |
| A4 | $1.0 \times 10^5$ | 0 | $5.0 \times 10^1$ | $1.0 \times 10^5$ | $8.0 \times 10^3$ | $3.3 \times 10^4$ |
| A5 | $1.0 \times 10^5$ | 0 | 0 | $2.0 \times 10^3$ | $1.0 \times 10^3$ | $1.0 \times 10^3$ |
| A6 | $3.7 \times 10^3$ | 0 | 0 | $1.5 \times 10^3$ | $4.7 \times 10^3$ | $7.2 \times 10^3$ |
| A10 | $3.0 \times 10^5$ | $1.1 \times 10^2$ | $1.3 \times 10^2$ | $1.0 \times 10^3$ | $1.2 \times 10^3$ | $1.5 \times 10^3$ |
| A16 | $8.0 \times 10^5$ | $1.0 \times 10^2$ | $1.2 \times 10^2$ | $1.0 \times 10^5$ | $1.0 \times 10^3$ | $1.6 \times 10^4$ |
| A19 | $1.2 \times 10^4$ | 0 | 0 | $1.0 \times 10^4$ | 0 | 0 |
| A22 | $3.5 \times 10^4$ | 0 | 0 | 0 | 0 | 0 |
| A23 | $8.0 \times 10^4$ | $1.3 \times 10^2$ | $1.2 \times 10^2$ | $1.0 \times 10^4$ | $1.0 \times 10^3$ | $1.0 \times 10^3$ |

\* See Table 1 for the origin of the samples.

### Propagation and purification of *Propionibacterium* phages

For phage purification, a single phage plaque was picked from the soft agar layer and added to 3 mL of an early-log-phase culture of the sensitive strain CIRM-BIA15 or CIRM-BIA509 (**Table 3**). The infected cultures were incubated at 30°C for 48 h. After incubation, the phage-infected cultures were filtered (0.2 µm) and tested again on the sensitive strain using YELA plates. This plaque-purification process was repeated for a total of three rounds for each phage. The resulting phage filtrates were stored at 4°C. Phage titer was first approximately determined by serial 10-fold dilutions of the filtrates. Then, each phage was amplified on plates by incubating 200 µL from a mid-log phase bacterial culture (OD_600_ = 0.4 – 0.7) with 100 µL of the diluted filtrate forming confluent lysis plaques for 20 min at 30°C, followed by plating using the soft agar method. For each phage, 10 plates were prepared, and 5 mL of YEL broth was added to each plate. After 20 min incubation at room temperature, the soft agar layer from multiple plates were recovered in several tubes and stored at 4°C overnight. The collected samples were then centrifuged at 9,000 x *g* for 20 min, and the supernatants were filtered (0.2 µm) and pooled. Lysates containing high-titer phages (10^9^-10^11^ PFU/mL) were further concentrated by ultracentrifugation at 35,000 x *g* for 2 h. The resulting pellets were gently resuspended in 1 mL of TM buffer (50 mM Tris-HCl, 10 mM MgSO_4_, pH 7.5). Phage particles were then purified from these suspensions by CsCl gradient ultracentrifugation (38) to obtain purified phage preparations at concentrations of 10^12^ - 10^13^ PFU/mL. The final phage stock suspensions were stored at 4°C or at -80°C with the addition of 15 % (vol/vol) glycerol.

**Table 3.** Main characteristics of phages isolated from Swiss-type cheeses.

| Phage name | Cheese sample | Propagation host | Nuclease S1 sensitivity | Phage morphology | Phage lenght (nm) | Phage width (nm) | Genome size (nucleotides) | GC content (%) | Number of predicted ORF | Genome accession number |
| --- | --- | --- | --- | --- | --- | --- | --- | --- | --- | --- |
| MINOG 1 | A3 | CIRM-BIA15 | Yes | Filamentous | 771 ± 34 | 11.9 ± 1.8 | 5,910 | 64.3 | 12 | ERZ24815863 |
| MINOG 2 | A4 | CIRM-BIA15 | Yes | Filamentous | 784 ± 39 | 11.8 ± 1.0 | 5,902 | 64.1 | 12 | ERZ24815864 |
| MINOG 3 | A5 | CIRM-BIA15 | Yes | Filamentous | 814 ± 73 | 11.1 ± 1.0 | 5,924 | 64.7 | 13 | ERZ24815865 |
| MINOG 4 | A6 | CIRM-BIA15 | Yes | Filamentous | 837 ± 41 | 11.6 ± 1.2 | 5,887 | 64.3 | 12 | ERZ24815866 |
| MINOG 5 | A16 | CIRM-BIA15 | Yes | Filamentous | 839 ± 61 | 12.2 ± 1.4 | 5,888 | 64.6 | 13 | ERZ24815867 |
| MINOG 6 | A10 | CIRM-BIA15 | Yes | Filamentous | 826 ± 40 | 11.3 ± 0.8 | 5,888 | 64.6 | 12 | ERZ24815868 |
| MINOG 7 | A19 | CIRM-BIA15 | Yes | Filamentous | 847 ± 80 | 12.6 ± 1.2 | 5,942 | 64.3 | 13 | ERZ24815869 |
| MINOG 8 | A22 | CIRM-BIA15 | Yes | Filamentous | 783 ± 42 | 10.2 ± 1.2 | 5,822 | 64.2 | 13 | ERZ24815870 |
| MINOG 9 | A23 | CIRM-BIA15 | Yes | Filamentous | 816 ± 43 | 10.3 ± 0.6 | 5,876 | 64.3 | 13 | ERZ24815871 |
| MINOG 10 | A5 | CIRM-BIA509 | No | Siphophage | ND | ND | ND | ND | ND | NA |
| MINOG 11 | A6 | CIRM-BIA509 | No | Siphophage | ND | ND | ND | ND | ND | NA |

ND, Not determined ; NA, Not applicable

### Transmission Electron Microscopy

A high-titer purified phage suspension (10^12^ - 10^13^ PFU/mL) was applied to freshly glow-discharged 200-mesh Formvar-coated copper grids and negatively stained with 2% uranyl acetate. The samples were examined at 120 kV with a JEOL 1400 transmission electron microscope equipped with an SC 1000 camera (Gatan Orius).

### Efficiency of plating

The efficiency of plating (EOP) for each phage was assessed using five strains of *P. freudenreichii.* To achieve this, 100 µL of overnight bacterial cultures were added to test tubes containing 3 mL of molten soft YEL agar (0.7% agar), which were mixed and overlaid on YELA plates. Next, 3 µL of a high-titer purified phage suspension (> 10^9^ PFU/mL) were spotted onto the agar plates containing the host strains. After incubation at 30°C for 3 days, plaques were enumerated on the plates.

### Phage DNA extraction and nuclease S1 treatment

*P. freudenreichii* phage genome was isolated from high-titer purified phage suspensions using the protocol developed for the isolation of ssDNA extraction from M13 phage using QIAGEN® Plasmid Kits. To confirm the presence of ssDNA, the isolated phage DNA (150-200 ng) was mixed with S1 nuclease (20 U, Promega), incubated at 37°C for 15 min, and analyzed on a 1% agarose gel.

### *Propionibacterium* phage genome sequencing and analysis

After extracting ssDNA from each phage, the synthesis of the second strand of complementary DNA was performed using the NebNext Second Strand Synthesis kit (BioLabs). A purification step using AMPure XP magnetic beads (Agencourt) was then performed. The libraries for DNA sequencing were subsequently prepared using the Nextera XT DNA Library Prep Illumina kit and sequenced during a paired-end run in 2x150bp. Sequencing read quality was assessed using FastQC 0.11.90 (39) before and after cleaning using PRINSEQ 0.20.4 (40) and Cutadapt 3.4 (41). The reads were subsampled before being assembled with SPAdes 3.15.4 (42). Assembly quality was assessed using Bowtie2 2.4.4 (43) and Samtools 1.12 (44). Structural and functional annotation was performed using Rime Bioinformatics’ rTOOLS2 pipeline. Genome comparison and alignments were performed using Clustal Omega - 1.2.4 (45). Annotated genomes of phages MINOG1 to MINOG9 have been deposited in the European Nucleotide Archive (ENA, Cambridge) under accession numbers ERZ24815863-ERZ24815871.

### Propionibacterium freudenreichii genome sequencing

Genomic DNA of strains CIRM-BIA15, CIRM-BIA129, CIRM-BIA508, CIRM-BIA1101, CIRM-BIA1102, CIRM-BIA1402, LSP100, and LSP102 was extracted from bacterial cultures using a phenol–chloroform protocol with enzymatic lysis. Briefly, 5 mL of an overnight culture (OD_600_ = 0.8 – 1.0) were centrifuged at 7,000 rpm for 5 min, and the resulting pellet was resuspended in 600 µL of buffer B1 (Qiagen) supplemented with RNase A (10 mg/mL, 10 µL, 100 µg) and lysozyme (50 mg/mL, 100 µL, 5 mg). The suspension was incubated at 42 °C for 30 min, followed by the addition of proteinase K (20 mg/mL, 50 µL, 1 mg) and a further incubation at 55 °C for 30 min to ensure complete cell lysis. An equal volume of phenol–chloroform–isoamyl alcohol (25:24:1, v/v/v) was added, mixed thoroughly, and centrifuged at 12,000 × *g* for 10 min to separate the phases. The aqueous phase containing genomic DNA was carefully transferred to a new tube, and DNA was precipitated by adding 0.7 volume of isopropanol, followed by centrifugation at 12,000 × *g* for 10 min. The resulting DNA pellets were washed with 70% ethanol, briefly air-dried to remove residual solvent, and resuspended in nuclease-free water according to standard procedures (46).

Genomes were sequenced by Illumina NovaSeq 2 x 150 pb and Oxford Nanopore (Eurofins Genomics, Constance, Germany). Genomes of strains CIRM-BIA15 and LSP102 were assembled with hybracter v0.11.2 (47) and genomes of strains CIRM-BIA129, CIRM-BIA508, CIRM-BIA1101, CIRM-BIA1102, CIRM-BIA1402, and LSP100 were assembled with Unicycler v0.5.1 (48). The quality of the genomes was assessed with QUAST v5.3.0 (49), Checkm2 v1.1.0 (50), Busco v.5.8.3 (51) and by aligning each genome against itself with Blastn v2.16.0 (52) to ensure the absence of large artifactual duplicated regions. Annotation was done with Bakta (53) and annotated genomes are available at the ENA (Cambridge) under accession number PRJEB109796.

### Bioinformatics analysis

Phages genomes were aligned and drawn with clinker (54). FastANI software (55) was used to compute pairwise ANI (Average Nucleotide Identity) values among phages genomes and the resulting heatmap was drawn with R package ggplot2. CRISPR arrays and their associated Cas systems were predicted from the genome sequences of *Propionibacterium freudenreichii* strains using CRISPRCasFinder platform (56, 57).

### Identification of *Propionibacterium* phage proteins using mass-spectrometry

To estimate the protein content of phage preparations, Qubit™ fluorometer (Invitrogen, Oregon, USA) measurements were carried out on 2 µL of purified phages. Depending on the result, between 10-15 µL of purified phage preparations (around 8 µg of protein per sample) were mixed with 2× Laemmli sample buffer and heated at 100 °C for 5 min. Then, the denatured proteins were loaded into each well of a 12 % SDS-PAGE gel (Miniprotean II, Bio-Rad). Electrophoresis was controlled by a pre-stained molecular mass markers (Precision Plus Protein™ Kaleidoscope™ Prestained Protein Standards, Bio-Rad) and stopped as soon as the yellow band of the size marker was visualized (about ½ hour). Gel pieces containing the protein samples were excised, and subjected to in-gel trypsinolysis followed by peptide extraction, as described previously (58). After digestion, the peptides were stored at -20°C until further analysis. Nano-liquid chromatography tandem mass spectrometry (LC-MS/MS) experiments were performed as previously described (59, 60). The peptides were identified from MS/MS spectra using i2MassChroQ software (61), and searches were performed against the genome sequence of filamenteous *Propionibacterium* phages, as described previously (58).

## RESULTS AND DISCUSSION

### Isolation of *Propionibacterium freudenreichii* phages from cheese samples

The presence of phages in thirty cheeses produced from either raw or heated milk (**Table 1**) was determined using ten *P. freudenreichii* strains as potential host strains. Phages infecting *P. freudenreichii* were found in nine out of the 30 cheeses (30%) (**Table 2**). All the phage-positive cheeses, sourced from different regions of France, shared the common feature of being made from raw milk and following Swiss-type cheese production method. Six out of the 10 strains were sensitive to some of these phages with titers ranging from 5.0 x 10^1^ (CIRM-BIA1101) to 8.0 x 10^5^ (CIRM-BIA15) PFU/g (**Table 2**). While some strains were sensitive to only a few phage samples (*e.g.*, CIRM-BIA129), others, such as CIRM-BIA15 and CIRM-BIA509 were susceptible to nearly all of them. Following purification, eleven phages were isolated using strains CIRM-BIA15 or CIRM-BIA509 as host, for further characterization (**Table 3**).

### Morphological characterization of *Propionibacterium freudenreichii* phages isolated from Swiss-type cheeses

The presence of phages was confirmed by examining the lysates using transmission electron microscopy. Nine of the eleven phages, designated MINOG1 to MINOG9, exhibited long, thin, flexible viral particles with uniform diameter and lacking distinct head–tail structures, characteristics of filamentous phages (**Figure 1A**, **Figure S1**). Their dimensions were nearly identical to those previously reported for the reference *Propionibacterium* filamentous phage B5 (**Table 3**) (30). The two other phages isolated, named MINOG10 and MINOG11, displayed morphological features typical of siphophages. These include an icosahedral capsid and a long non-contractile tail with a terminal baseplate (**Figure 1A**), a morphology that represents the majority of phages described to date infecting *P. freudenreichii* (25–28).

**Figure 1.**
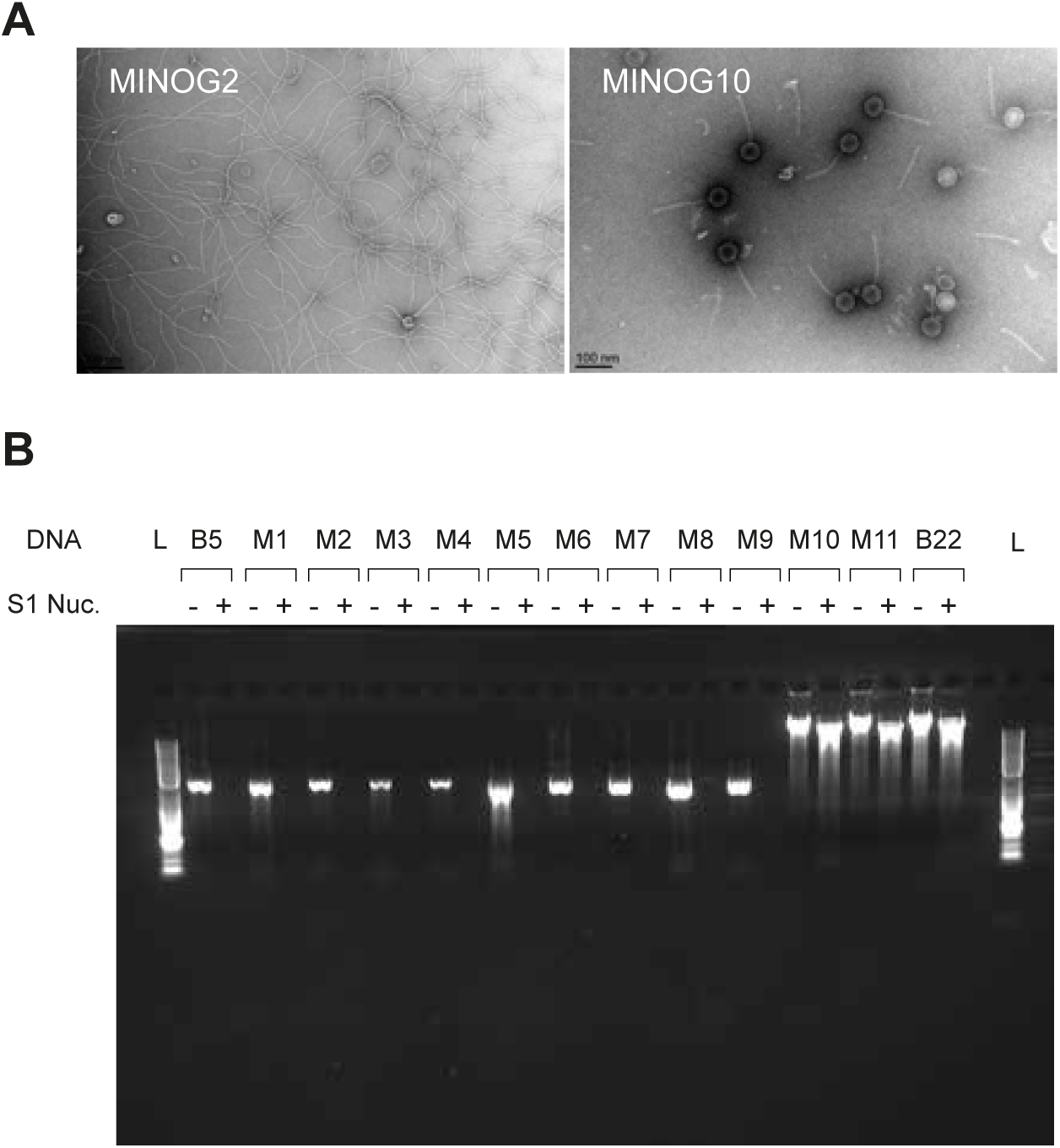
Characterization of *Propionibacterium* phages isolated from Swiss-type cheeses. (A) Electron micrographs of the filamentous phage MINOG2 as well as the siphovirus MINOG10. (B) Activity of S1 nuclease on phage DNA samples. L, DNA molecular weight marker (SmartLadder, Eurogentec); B5, single-stranded DNA from filamentous *Propionibacterium* phage B5 used as positive control; M1 to M11, DNAs of *Propionibacterium* phages MINOG1 to MINOG11; B22, double-stranded DNA from *Propionibacterium* phage B22 used as negative control; - and +, S1 nuclease untreated and treated DNA samples, respectively.

Given the scarce descriptions of filamentous phages infecting Gram-positive bacteria, we undertook the characterization of the nine newly isolated putative filamentous phages. To first validate their classification as filamentous phages, which typically harbor ssDNA genomes, a S1 nuclease protection assay was carried out on the extracted phage genomes (**Figure 1B**, **Table 3**). DNA from the filamentous phage B5 (ssDNA) and the siphophage B22 (dsDNA), both infecting *P. freudenreichii,* were used as controls (Gautier et al. 1992; Chopin et al. 2002). As expected, DNA from the two new siphophages were resistant to S1 nuclease treatment, supporting the presence of dsDNA genomes (**Table 3**). In contrast, the DNA of the nine phages exhibiting filamentous morphologies was degraded by the S1 nuclease, indicating that their genomes consist of ssDNA, as observed for phage B5 (**Figure 1B**, **Table 3**). Thus, nine filamentous phages infecting *P. freudenreichii* were isolated from various Swiss-type cheeses produced at different manufacturing sites in France.

### Host range of Propionibacterium freudenreichii filamentous phages

To assess the host range of the newly isolated *Propionibacterium* filamentous phages, the efficiency of plating (EOP) of each phage was tested on five *P. freudenreichii* indicator strains (**Figure 2**). The strain CIRM-BIA15 was chosen as the primary host due to its highest sensitivity among all tested phages. Based on their infection patterns, the nine new filamentous phages and the control phage B5 were classified into four groups. The first two groups include phages that exhibited low productive infections on strains CIRM-BIA129 and CIRM-BIA1101 (EOP < 0.001) but differed on their infectivity on strains LSP100 and LSP102. Group I phages (B5, MINOG2, and MINOG9) demonstrated high productive infection (EOP > 0.1) against strains LSP100 and LSP102, whereas group II phages (MINOG1, MINOG6 and MINOG7) showed only moderate productive infection (0.001 < EOP < 0.1). Group III comprised phages MINOG3, MINOG4, and MINOG8, which exhibited low productive infection across all tested strains (EOP < 0.00099). Finally, MINOG5, the unique member of group IV, displayed moderate productive infection across all tested strains (0.001 < EOP < 0.1). This analysis partly supports our previous findings on phage titer from cheese samples, showing the broader host range and infectivity of phages MINOG2, MINOG5, and MINOG9 (**Table 2**). Although the phages from cheese samples A10, A16, and A23 had similar host ranges, only the phages isolated from cheeses A16 and A23 (*i.e.*, MINOG5 and MINOG9) exhibited a broad host range, suggesting that cheese A10 contained additional phages beyond MINOG6. Overall, our results indicate that *P. freudenreichii* filamentous phages exhibit a diverse host range.

**Figure 2.**
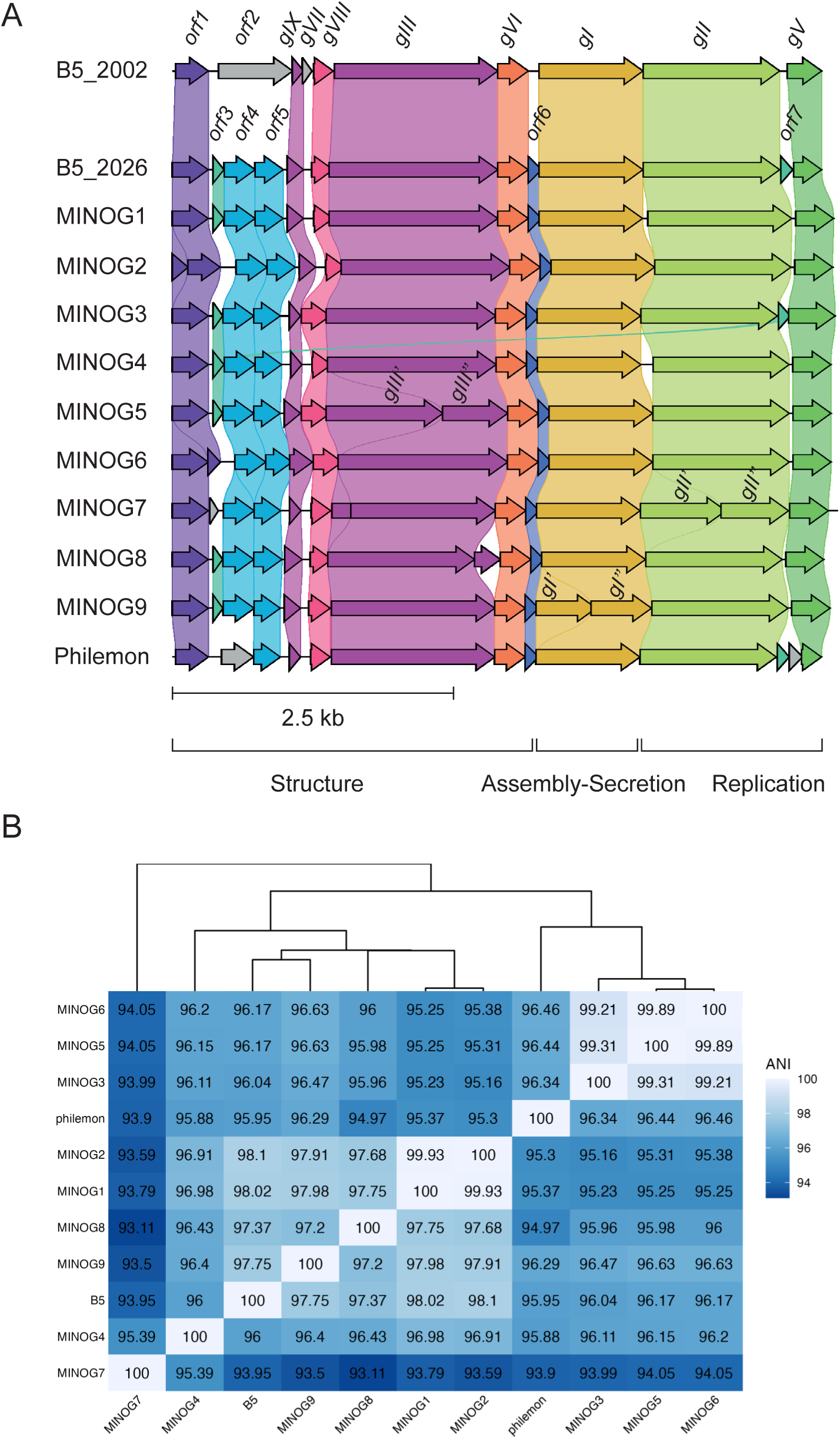
Genomic and phylogenomic comparison of *P. freudenreichii* filamentous phages. (A) Comparison of the genomic organization of *P. freudenreichii* filamentous phages. B5_2002 corresponds to the original annotation of the phage B5 genome, first established in 2001 (30). B5_2026 designates the reanalysis of the phage B5 genome performed in this study. Each group of homologous genes is represented by the same color. The functional modules for structure, assembly-secretion, and replication are indicated according to the Ff phage model proposed by Chopin et *al.* (30). The *gVIII* gene encodes the major coat protein pVIII (also designated CoaB). The *gIII*, *gVI, gVII*, and *gIX* genes encode the minor coat proteins pIII (also designated CoaA), pVI, pVII, and pIX, respectively. The *gI*, *gII*, and *gV* genes encodes the morphogenesis protein pI, DNA rolling circle replication protein pII, and single-stranded DNA binding protein pV, respectively. The functions of the proteins encoded by the others ORFs remain unknown. (B) Comparison of the average nucleotide identity (ANI) among the *Propionibacterium* filamentous phages. The heatmap showed the pairwise percentage of identity between the genomes of eleven filamentous phages.

### Characterization of *Propionibacterium freudenreichii* filamentous phage genomes

To explore the genomic diversity of the newly isolated filamentous phages, we performed whole-genome sequencing and compared them with the genome of the reference phages B5 and Philemon. The genomes of the newly isolated filamentous phages ranged in size from 5,822 bases (MINOG8) to 5,942 bases (MINOG7), with an average GC content of 64.37% (**Table 3**). The genome of B5 and Philemon are slightly smaller with 5,802 and 5,806 bases, respectively. A total of 12 to 13 open reading frames (ORFs) were predicted from the complete genome of these isolated filamentous phages, most of which had no detectable homologs in public databases, except for those found in B5 and Philemon (**Figure 2A**). To facilitate comparison, we used the ORF annotation system proposed by (30), which is also based on the gene annotations of Ff phages, *i.e.*, M13, f1, and Fd (62, 63), the archetypal filamentous phages that infects *Escherichia coli*, with genes designated as *gI* to *gXIII* and the corresponding proteins named pI to pXIII. When no homologous genes could be identified, annotations were complemented using ORF numbering.

The number of detected ORFs exceeded the 10 ORFs previously reported for phage B5, which was annotated more than 20 years ago using earlier bioinformatic tools. Reannotation of B5 genome resulted in an ORF content similar to that observed in the newly identified *Propionibacterium* filamentous phages, with most original ORFs retained between the two annotation versions (**Figure 2A**). Nevertheless, some differences were observed. Notably, the ORF encoding the pVII homolog of Ff phage, a minor capsid protein, was not identified in the new automatic annotation version of the B5 genome, although a corresponding ORF could be detected manually. In addition, the region corresponding to *orf2* in the original annotation of B5, which was predicted as a single ORF, was resolved into three distinct ORFs in the new annotation (*orf3, orf4, orf5*).

The *Propionibacterium* filamentous phage genomes exhibited collinear gene organization on the same strand (**Figure 2A**). Moreover, as observed for filamentous phages infecting Gram-negative bacteria, genes are organized into three functional modules: i) structural, ii) assembly and secretion, and iii) replication (30, 63). The structural module notably includes genes coding for the major coat protein pVIII, which forms the body of the filamentous particles (*e.g.*, CoaB), and helically wraps around the phage ssDNA, as well as the minor coat proteins pIII (*e.g.*, CoaA), pVI, pVII, and pIX, which play key roles in particle assembly, stability, and infectivity (62, 64, 65). The assembly and secretion module appears to comprise of a single gene encoding the morphogenesis protein pI, an ATPase of the FtsK-HerA superfamily that plays an essential role in the assembly of Gram-negative filamentous phages (30, 36, 66, 67). Note that an additional gene, absent from *Propionibacterium* filamentous phage genomes and encoding the outer membrane protein pIV, constitutes the assembly and secretion module in filamentous phages infecting Gram-negative bacteria (68–70). Finally, the replication module includes, notably, two genes, *gII* and *gV*, encoding a phage DNA rolling circle replication control protein (pII) and a ssDNA binding protein, functionally equivalent to pV that prevents complementary DNA strand synthesis (62, 71). The newly isolated phages display genomic features typical of filamentous phages.

### Identification of *Propionibacterium freudenreichii* filamentous phage proteins by mass spectrometry

As indicated above, the number of ORF detected and their annotations may vary depending on the analysis tools used. To validate the annotation of the filamentous phage structural proteins, the nine newly isolated phages and phage B5, were analyzed using LC-MS/MS. The proteomic analysis revealed the presence of five predicted protein-coding ORFs. Depending on the phage samples, between two (B5, MINOG8) and five (MINOG1, MINOG2) annotated proteins, primarily associated with the structural module, were detected (**Table 4**). Proteins pIII and pVIII were the most abundant and were detected in all samples. It has been estimated that each filamentous phage contains approximately 2,700 copies of pVIII, one of the most abundant phage proteins, that form the body of the phage, as well as 5 to 10 copies of minor coat proteins, such as pIII which plays a crucial role in the initial steps of infection. The third most abundant protein, ORF1, was detected in 7 out of 10 phage preparations. This protein is classified as hypothetical, with no known homologs in public database, including PhagesDB. Finally, the last two identified proteins were pI (assembly and export module) and pII (replication module), detected in only three and two phage preparations, respectively.

**Table 4.**
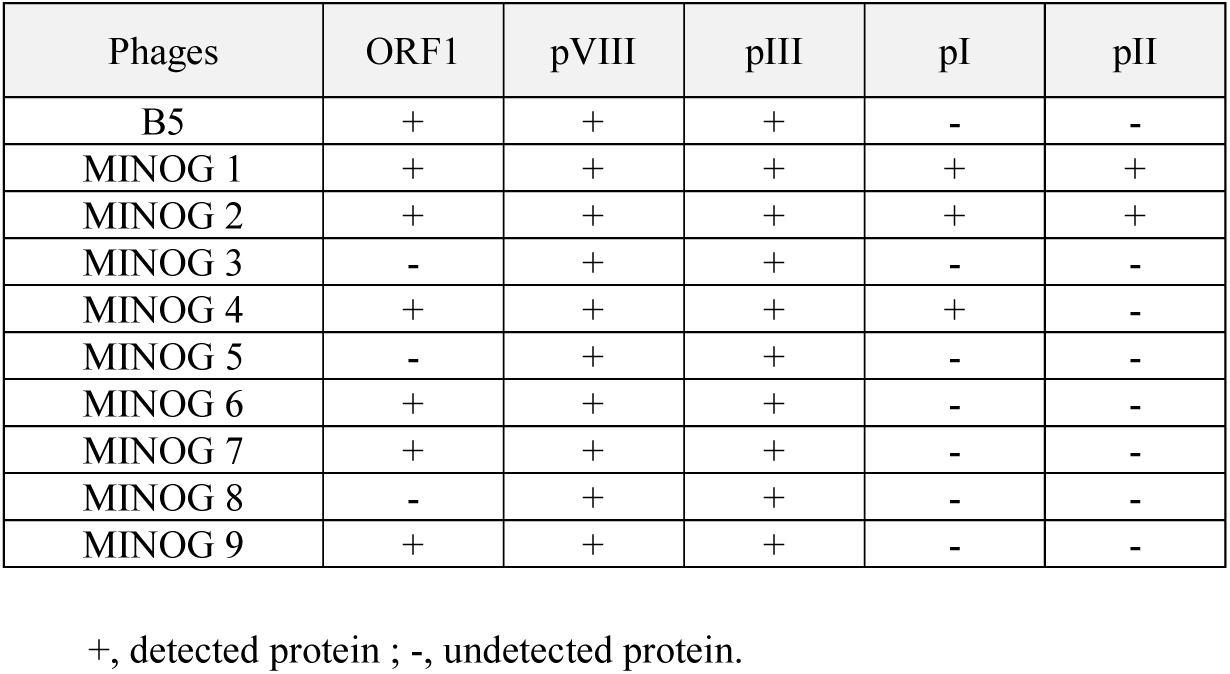
Identification of viral proteins by mass spectrometry according to the phages analyzed.

### Phylogenetic analysis of *Propionibacterium freudenreichii* filamentous phages

A pairwise comparison of all *P. freudenreichii* phage genomes, including B5 and Philemon, was performed using average nucleotide identity (ANI) (**Figure 3B**). The phages clustered into three groups. The first group consisted of six phages (B5, MINOG1, MINOG2, MINOG4, MINOG8, and MINOG9) with an ANI value between 94.97 and 99.93%. Notably, this group includes two nearly identical phages, MINOG1 and MINOG2, which shared 99.93% identity. The second groups comprised MINOG3, MINOG5, and MINOG6, all exhibiting identity greater than 99%. The phage Philemon was closely related to this group, with approximately 96% identity. The third group comprised only one phage (MINOG7) and was the most distant, with an identity ranging from 93.11 to 95.39% compared to the other phages. It is also noteworthy that phages MINOG5, MINOG7, and MINOG8, which belong to the three phylogenomic groups described above, were all isolated from the same cheese type (*i.e*., Comté) produced in the same French department *(i.e*., Doubs), further indicating that genomic diversity is not strictly linked to cheese type or production region.

**Figure 3.**
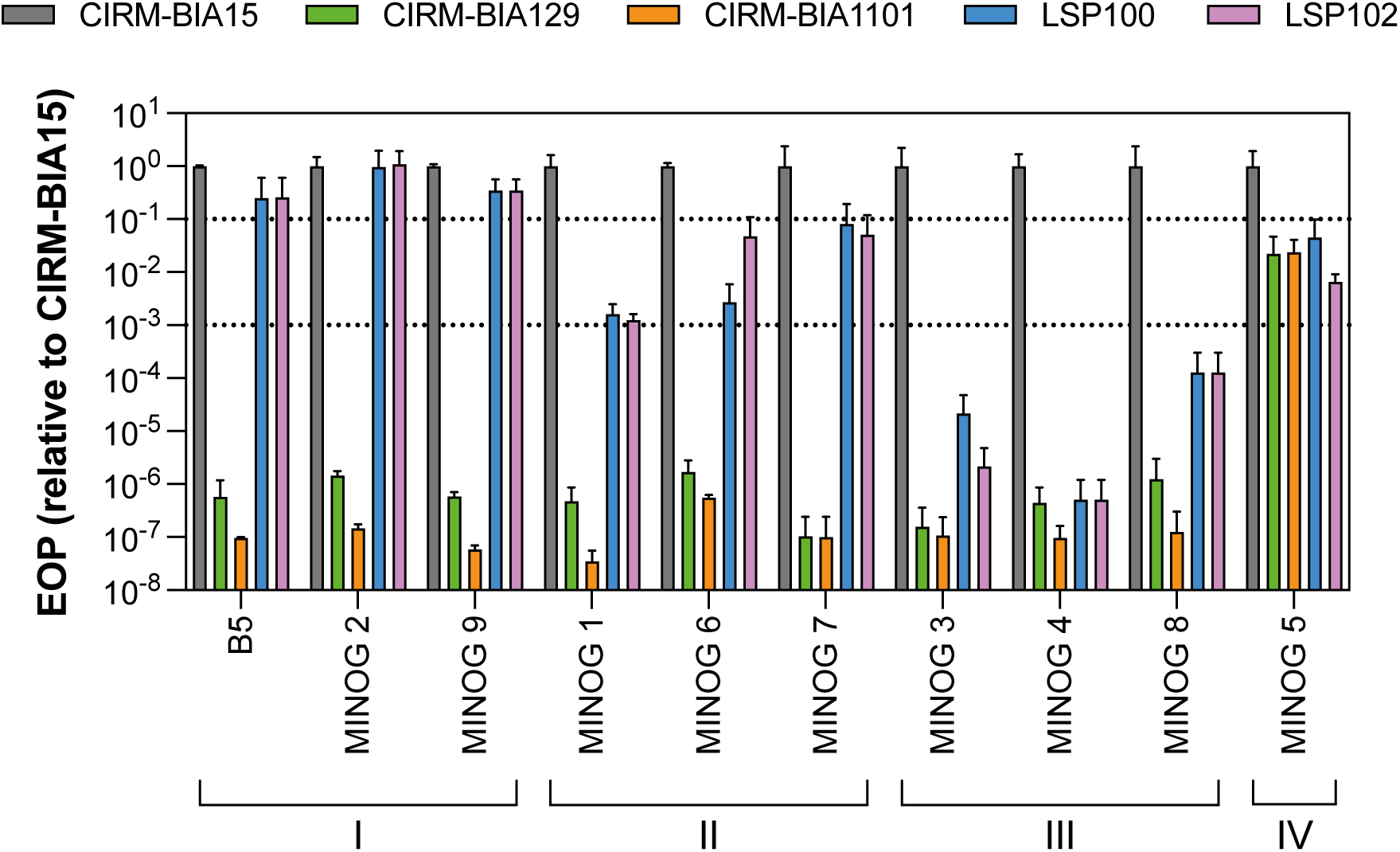
Efficiency of plating (EOP) of *Propionibacterium* filamentous phages on *P. freudenreichii* strains. The EOP was calculated as the ratio of the average PFU obtained on the tested bacteria to that obtained on the host strain CIRM-BIA15. The standard deviation was determined from two independent experiments performed in duplicate. Phages were clustered in four groups, from I to IV, based on their pattern of infection.

Overall, this comparative genomic analysis highlighted strong genetic conservation within closely related groups but with notable genomic diversity across the *P. freudenreichii* filamentous phage population. Moreover, this analysis showed that host range groupings (**Figure 2**) do not fully match phylogeny (**Figure 3B**). In particular, closely phylogenetically related phages, *i.e*., MINOG5 and MINOG6, which shared 99.89% sequence identity, displayed different host range, highlighting that minor genomic variations may substantially influence phage–host interactions.

### CRISPR spacer analysis in *Propionibacterium freudenreichii* genomes and evidence of past phage exposure

A previous study indicated that most *P. freudenreichii* strains contain CRISPR-Cas systems (types I-G and I-E) (72), presumably acting as a defense system against phages (73). The genome sequences of eight *P. freudenreichii* strains (CIRM-BIA15, CIRM-BIA129, CIRM-BIA508, CIRM-BIA1101, CIRM-BIA1102, CIRM-BIA1402, LSP100, and LSP102) used in this study were screened for CRISPR spacers targeting *Propionibacterium* filamentous phages. High-confidence CRISPR arrays and associated Cas systems were identified in the genomes of only six strains (**Table S1**). The genomes harbored between one to four CRISPR arrays, comprising 9 to 89 spacers. Among them, the genomes of strains LSP100 and LSP102 contained a single identical 38-nucleotide spacer within a predicted CRIPSR array, showing high sequence identity to all *Propionibacterium* filamentous phages (**Table 5** and **Table S3)**. This sequence exhibited from 79 (MINOG9) to 97% (MINOG7) identity with a phage genomic region, depending on the phage. The corresponding potential protospacers were in the *gV* gene coding for the pV ssDNA binding protein.

**Table 5.**
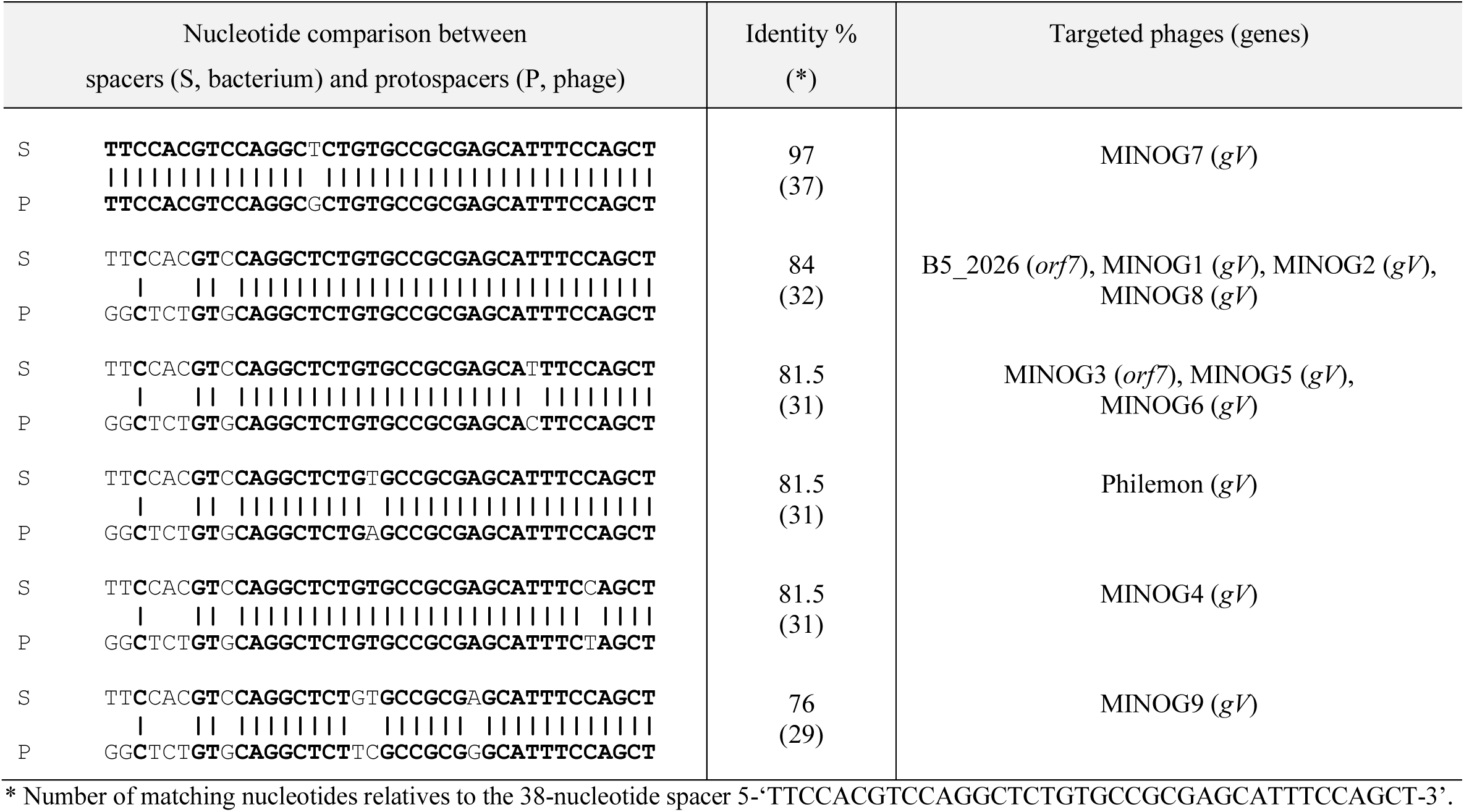
Homology regions between an identified CRISPR spacer in the genomes of strains *P. freudenreichii* LS 2500 and 2502 and protospacers in filamentous phage genomes.

To determine whether other *P. freudenreichii* strains possess spacers targeting the eleven *Propionibacterium* filamentous phage genomes, we analyzed the 136 NCBI *P. freudenreichii* genomes using Blastn (release June 2026). The genomes of eleven *P. freudenreichii* strains harbored CRISPR loci containing spacers ranging from 33 to 38 nucleotides in length and showing high sequence homology (from 79 to 100% identity) to the 11 *Propionibacterium* filamentous phage genomes (**Table S2** and **Table S3**). These genomes also comprised between one and four CRISPR arrays, all associated with Type I CRISPR-Cas systems, and containing 9 to 113 spacers. Among the 823 spacers identified across these strains, including LSP100 and LSP102, thirteen (1.6%) showed sequence identity to genomic region of *Propionibacterium* filamentous phage (**Table S3**). Four identical spacers were found in up to three different strains. While the genomes of most strains (11/13) harbored only a single spacer targeting these filamentous phages, strains FAM23877 and PFRJS9 contained two and six different spacers against them, respectively, suggesting more extensive exposure to these phages.

Of the eleven filamentous phages analyzed, seven contained protospacers in their genomes matching the 13 identified spacers. Amongst the exceptions, phage MINOG7 harbored only 11 protospacers, whereas MINOG4 contained 14 due to a protospacer duplication. The identified protospacers mapped to genes *gII*, *gIII*, *orf1*, *orf12*, *orf15*, *gI, orf16, and gV5,* with *gII* and *gIII* being the most targeted. It should be noted that type I CRISPR-Cas systems have been shown previously to acquire spacers from the ssDNA filamentous coliphage M13 (74). Moreover, phage-matching spacers can even stimulate the acquisition of more M13-derived spacers, a phenomenon known as priming. Altogether, our findings clearly indicate that *P. freudenreichii* strains are exposed to filamentous phages isolated from cheeses, even when the strains were isolated from non-dairy environments. For example, strain P.UF1was isolated from the feces of a human breast milk-fed preterm infant (75).

## CONCLUSION

We isolated and characterized nine novel filamentous phages infecting the Gram-positive bacterium *P. freudenreichii*. To our knowledge, this represents only the second demonstration of filamentous phages infecting Gram-positive bacteria. Comparison with the two previously characterized *P. freudenreichii* filamentous phages reveals a conserved genetic organization and gene repertoire. While their genome varied only from 0.1 to 7%, they exhibited distinct host range specificities. No clear relationship was identified between isolation source, genetic diversity, host range, and CRISPR spacer in *P. freudenreichii* genomes. Several lines of evidence suggest additional defense systems beyond CRISPR-Cas in *P. freudenreichii*. Strains CIRM-BIA129 and CIRM-BIA1101 were resistant to nearly all tested phages despite lacking CRISPR spacers targeting the genome of these phages. Likewise, *P. freudenreichii* strains LSP100 and LSP102 showed different phage sensitive profiles despite exact spacer-protospacer matches with phages B5, MINOG1, MINOG2, and MINOG8. The factors underlying these phage–host interactions warrant further investigation and could provide insights into the mechanisms bacteria use to defend against ssDNA phages.

## Acknowledgments

The authors warmly thank the Laboratoires STANDA (https://www.standa-fr.com), notably Christophe Hervé and Riwanon Lemé, for providing strains LSP100 and LSP102. We are grateful to the INRAE MIGALE bioinformatics facility for providing help, computing and storage resources(76). S.M. acknowledges funding from the Natural Sciences and Engineering Research Council of Canada (Discovery Program) and the FRQNT strategic network Op+Lait. S.M. held the Canada Research Chair in Bacteriophages during part of the study.

